# Differential excitability of PV and SST neurons results in distinct functional roles in inhibition stabilization of Up-states

**DOI:** 10.1101/2020.11.26.395343

**Authors:** Juan L. Romero-Sosa, Helen Motanis, Dean V. Buonomano

**Affiliations:** Department of Neurobiology, Integrative Center for Learning and Memory, University of California, Los Angeles, Los Angeles, 90095; Department of Psychology, University of California, Los Angeles, Los Angeles, CA. 90095; Department of Neurosurgery, University of California, Los Angeles, Los Angeles, CA. 90095

**Keywords:** Up-states, neural dynamics, inhibition, parvalbumin, somatostatin, neural network model

## Abstract

Up-states are the best-studied example of an emergent neural dynamic regime. Computational models based on a single class of inhibitory neurons indicate that Up-states reflect bistable dynamical systems in which positive feedback is stabilized by strong inhibition and predict a paradoxical effect in which increased drive to inhibitory neurons results in decreased inhibitory activity. To date, however, computational models have not incorporated empirically defined properties of PV and SST neurons. Here we first, experimentally characterized the frequency-current (F-I) curves of pyramidal, PV, and SST neurons and confirmed a sharp difference between the threshold and slopes of PV and SST neurons. The empirically defined F-I curves were incorporated into a three-population computational model that simulated the empirically-derived firing rates of pyramidal, PV, and SST neurons. Simulations revealed that the intrinsic properties were sufficient to predict that PV neurons are primarily responsible for generating the nontrivial fixed points representing Up-states. Simulations and analytical methods demonstrated that while the paradoxical effect is not obligatory in a model with two classes of inhibitory neurons, it is present in most regimes. Finally, experimental tests validated predictions of the model that the Pyr↔PV inhibitory loop is stronger than the Pyr↔SST loop.

**SIGNIFICANCE STATEMENT:** Many cortical computations, such as working memory, rely on the local recurrent excitatory connections that define cortical circuit motifs. Up-states are among the simplest and best studied examples of neural dynamic regimes that rely on recurrent excitatory excitation. However, this positive feedback must be held in check by inhibition. To address the relative contribution of PV and SST neurons we characterized the intrinsic input-output differences between these classes of inhibitory neurons, and using experimental and theoretical methods show that the higher threshold and gain of PV leads to a dominant role in network stabilization.

## INTRODUCTION

Many neural computations emerge from the intrinsic dynamics generated by the recurrent connectivity of neocortical microcircuits (Douglas and Martin, 2007; Rabinovich et al., 2008; Buonomano and Maass, 2009; Chaisangmongkon et al., 2017). Such intrinsically generated dynamic regimes are hypothesized to underlie a wide range of neural computations, including those in which the brain must temporarily store information (e.g., working memory) or produce appropriately timed responses in the absence of ongoing external inputs (Hebb, 1949; Hopfield, 1982; Wang, 2001; Mauk and Buonomano, 2004; Huertas et al., 2015). Indeed, *in vivo* studies have implicated internally generated dynamics in a range of memory and temporal computations (Quintana and Fuster, 1992; Funahashi et al., 1993; Shuler and Bear, 2006; Carnevale et al., 2015; Namboodiri et al., 2015; Guo et al., 2017; Inagaki et al., 2019). Some of these dynamic regimes, including self-sustained tonic and dynamically changing patterns of activity, have also been observed in *in vitro* and ex *vivo* circuits (Sanchez-Vives and McCormick, 2000; Wang, 2001; Johnson and Buonomano, 2007; Mann et al., 2009; Sadovsky and MacLean, 2014; Carnevale et al., 2015; Carrillo-Reid et al., 2015; Dechery and MacLean, 2017), suggesting that the learning rules underlying internally generated neural dynamics are local and operational ex vivo. Yet, while it is well established that emergent dynamic regimes rely on the presence of recurrent excitation and balanced excitation and inhibition, relatively little is known about how self-sustaining dynamics emerge in cortical circuits in a self-organizing manner.

The best-studied form of emergent dynamics in neocortical circuits are Up-states, a term that generally refers to network-wide regimes in which excitatory neurons can transition between quiescent Down-states to more-or-less stable depolarized states with low to moderate firing rates (Sanchez-Vives and McCormick, 2000; Neske et al., 2015; Bartram et al., 2017). Up-states can occur spontaneously or be evoked and are observed *in vivo* during anesthesia, slow-wave sleep, quiet wakefulness (Steriade et al., 1993; Timofeev et al., 2000; Beltramo et al., 2013; Hromádka et al., 2013), and in acute slices (Sanchez-Vives and McCormick, 2000; Shu et al., 2003; Fanselow and Connors, 2010; Sippy and Yuste, 2013; Xu et al., 2013; Sadovsky and MacLean, 2014; Neske et al., 2015; Bartram et al., 2017), as well as in ex *vivo* cortical cultures (Plenz and Kitai, 1998; Seamans et al., 2003; Johnson and Buonomano, 2007; Kroener et al., 2009; Motanis and Buonomano, 2015; Motanis and Buonomano, 2020). Up-states have been proposed to have multiple functional roles, including memory consolidation and synaptic homeostasis (Tononi and Cirelli, 2003; Marshall et al., 2006; Sirota and Buzsáki, 2007; Vyazovskiy et al., 2008; Diekelmann and Born, 2010). It has also been hypothesized that Upstates are equivalent to the desynchronized regimes of awake cortex (Destexhe et al., 2007). For example, the voltage distribution of Up-states in awake cortex are indistinguishable from those observed during anesthesia (Constantinople and Bruno, 2011), and active sensory processing can shift cortical circuits to depolarized Up-state like patterns (Crochet and Petersen, 2006; Haider et al., 2007; Tan et al., 2014). The notion that Up-states are related to active cortical processing regimes is further reinforced by theoretical studies in which Up-states mirror so-called asynchronous regimes, wherein recurrently connected neurons are in a tonic, depolarized regime and spikes are triggered by ongoing fluctuations (Brunel, 2000; Renart et al., 2010; Tartaglia and Brunel, 2017). Together these studies suggest that Up-states reflect an important neural dynamic regime, and that neocortical circuits are programmed to seek out this regime under a wide range of conditions—including *in vivo* and ex *vivo* conditions.

Computational models have proposed that Up-states reflect intrinsically generated dynamic regimes in which self-sustaining activity is maintained through positive feedback and mediated through recurrent excitatory connections. This positive feedback is, in turn, held in check through rapid and strong inhibition, resulting in an *inhibition-stabilized network* (Tsodyks et al., 1997; Ozeki et al., 2009; Rubin et al., 2015; Jercog et al., 2017). To date, these models have effectively captured many aspects of Up/Down-state transitions and asynchronous network dynamics (Brunel, 2000; Holcman and Tsodyks, 2006; Renart et al., 2010; Dao Duc et al., 2015; Jercog et al., 2017; Tartaglia and Brunel, 2017). However, these models have focused primarily on a single unspecified class of inhibitory neurons. Yet it is well established that multiple classes of inhibitory neurons are active during Up-states recorded *in vivo* and in acute slices (Neske et al., 2015; Urban-Ciecko et al., 2015; Neske and Connors, 2016; Zucca et al., 2017). Thus, the respective functional roles of these interneurons remain mostly unaddressed.

Here we use experimental and computational methods to study the role of the two most common populations of inhibitory neurons—parvalbumin (PV) and somatostatin-positive (SST) neurons (Rudy et al., 2011)—in emergent dynamics. Experiments were performed in *ex vivo* organotypic cortical cultures to ensure that the experimental observations parallel our “standalone” computational model. Organotypic slices are a standard preparation to study synaptic plasticity and cortical microcircuit functions (Debanne et al., 1994; Hayashi et al., 2000; Esteban et al., 2003; Seamans et al., 2003; Kerr and Plenz, 2004; Goold and Nicoll, 2010; Yamada et al., 2010). *Ex vivo* organotypic slices provide a manner to unambiguously ascertain that the observed dynamics emerge locally within the circuit being studied. Additionally, consistent with the notion that Up-states reflect a core dynamic regime that cortical circuits are programmed to seek out, spontaneous and evoked Up-states emerge in cortical organotypic cultures over the course of *ex vivo* development (Johnson and Buonomano, 2007; Motanis and Buonomano, 2015; Motanis and Buonomano, 2020). Overall, our approach allowed us to directly compare experimental and model parameters of PV and SST neurons. Surprisingly, they reveal that the intrinsic properties of PV and SST neurons are sufficient to predict robust differential contributions of these inhibitory neuron classes emergent dynamics.

## METHODS

### Electrophysiology and ex vivo slices

All animal experiments followed guidelines established by the National Institutes of Health (NIH) and were approved by the Chancellor’s Animal Research Committee at the University of California, Los Angeles. Organotypic slices were prepared using the interface method (Stoppini et al., 1991; Buonomano, 2003). Five to seven-day-old PV-Cre and SST-Cre mice pups (Jackson Laboratory #017320 and #013044, respectively) were anesthetized with isoflurane and decapitated. The brain was removed and placed in chilled cutting media. Coronal slices (400 μm thickness) containing primary somatosensory and auditory cortex were sliced using a vibratome (Leica VT1200) and placed on filters (MillicellCM, Millipore, Billerica, MA, USA) with 1 mL of culture media. Culture media was changed at 1 and 24 hours after cutting and every 2-3 days thereafter. Cutting media consisted of EMEM (MediaTech cat. #15-010) plus (final concentration in mM): MgCl_2_, 3; glucose, 10; HEPES, 25; and T ris-base, 10. Culture media consisted of EMEM plus (final concentration in mM): glutamine, 1; CaCl_2_, 2.6; MgSO_4_, 2.6; glucose, 30; HEPES, 30; ascorbic acid, 0.5; 20% horse serum, 10 units/L penicillin, and 10 μg/L streptomycin. Slices were incubated in 5% CO2 at 35°C.

Recordings were performed at 23-35 days-in-vitro (DIV) in artificial cerebrospinal fluid (ACSF) composed of (in mM): 125 NaCl, 5.1 KCl, 2.6 MgSO_4_, 26.1 NaHCO_3_, 1 NaH_2_PO_4_, 25 glucose, and 2.6 CaCl_2_ (ACSF was formulated to match the standard culture media). The temperature was maintained at 32-33 °C, and the perfusion rate was set at 4.5-5 mL/min.

Recordings from pyramidal neurons relied on visualized whole-cell patching and were identified based on their intrinsic electrophysiological properties. Florescent PV and SST neurons were targeted using a 430 nm LED to visualize tdTomato. The internal solution for whole-cell recordings contained (in mM): 100 K-gluconate, 20 KCl, 4 ATP-Mg,10 phosphocreatine, 0.03 GTP-Na, and 10 HEPES and was adjusted to pH 7.3 and 300 mOsm. In order to be considered for analysis, cells had to have a resting potential less than −55 mV and not change by more than 15% over the course of the recording. The criteria for input and series resistance was 100-300 MΩ and <25 MΩ, respectively.

To estimate the average Up-state membrane potential during light stimulation and after light stimulation, the entire Up-state trace was filtered with a 10 ms median filter. Two 100 ms long windows were isolated from the trace, one during light stimulation and the other 100 ms after the end of light stimulation. These windows were averaged to get one membrane potential value per neuron.

### Transfection and Optogenetics

PV-Cre and SST-Cre slices were transfected in three different configurations: 1) 2.2 μL of ChETA (pAAV9-Ef1a-DIO-ChETA-EYFP Addgene#: 26968) and 1 μL of LSL-tdTomato (Addgene#: 100048); 2) 2.2 μL of DIO eNpHR 3.0 and 1 μL of LSL-tdTomato; or 3) 1 μL of LSL-tdTomato. The approximate titer of all viral solutions was 1 ×10^13^ vg/mL. Note that some of the paired recording data was obtained in slices transfected with halorhodopsin as part of other experiments. Transfection was performed at 5-7 DIV by gently delivering the virus mixture in a patch electrode at 3-5 positions in the cortex of the slice in order to transfect as many Cre-positive cells as possible. Experiments were performed at least 16-18 days after transfection.

Light stimulation was performed in a closed-loop fashion. Costume written code in Signal (Cambridge Electronic Design Limited) detected threshold crossings (approximately 5 mV) that marked the potential onset of an Up-state, which triggered light stimulation after a delay of 250 ms. Light stimulation consisted of either 15 pulses at 66 Hz (5 ms on) or 25 pulses at 50 Hz (10 ms on). Light stimulation was delivered via a royal blue (457 nm) LED (super-bright LEDs) at an intensity of 78 mW/cm^2^.

### Statistics

Non-repeated one-way ANOVA’s were also performed in MATLAB. The Up-state duration and voltage (Fig 7) were not normally distributed *(kstest* in MATLAB), thus we used paired nonparametric statistics to contrast these measures *(signrank* in MATLAB).

### Computational Model

Our three-population model was based on the two-population model of Jercog et al (2017). The model was composed of three classes of neurons representing the excitatory pyramidal neurons (*E*), PV neurons (*P*), and SST (*S*). Each of these populations was modeled as a single “unit” according to:

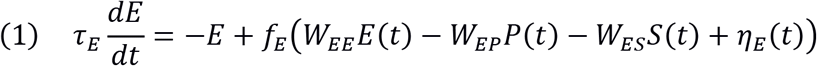

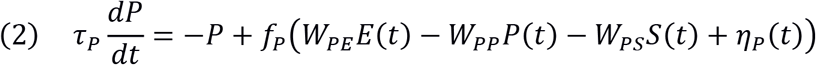

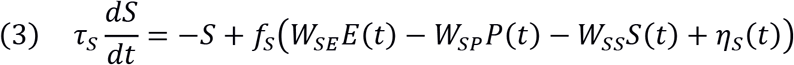

where *W_XY_* represents the weight between the postsynaptic unit *X* (*E, P,* or *S*) and presynaptic unit *Y* (*E, P*, or *S*). *τ_X_* and *η_X_* represent a time constant and an independent noise term, respectively. Similar to Jercog et al (2017), the noise term was an Ornstein-Uhlenbeck process with a time constant of 1 ms and a standard-deviation of σ_x_. We have not included an adaptation term that contributes to Up→Down because we focus primarily on the steady-state values. Thus, all the Up→Down and Down→Up transitions shown in the figures where fluctuation induced. Because all analyses focused on fixed-point Up-state values the results presented are independent of Up↔Down transitions. The function *f_Y_*(*x*) represents the intrinsic excitability of the three neuron classes, i.e., the F-I curve or activation function, which was simulated as a threshold-linear function characterized by a threshold (*θ_X_*) and gain that corresponds to the slope of the F-I curve (*g_X_*):

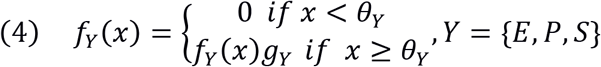

#### Paradoxical effect

To examine the paradoxical effect, we stimulated the *P* or *S* neurons by emulating an optogenetic experiment. Mathematically an external current is equivalent to decreasing the threshold of the F-I function (Eq. 4). Specifically, the threshold in the presence of simulated light activation can be written as: 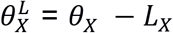, where the subscript *X* represents the *P* or *S* population, and *L_X_* captures the optogenetic activation of either population (*L_X_*=5 and 20 for the weak and strong activation, respectively).

### Empirical fits of the F-I functions

During whole-cell recordings, intrinsic excitability was measured with 250 ms current steps in the range of 0.05 to 0.6 nA depending on cell class. For each neuronal class, we fit the mean spike frequency x intensity (F-I) curve to a threshold-linear activation function (**Eq. 4**), which led to values threshold values of 0.06, 0.36, and 0.18 nA, and gains of 124, 334, 198 Hz/nA, for the Pyr, PV, and SST neuron classes, respectively. To simplify the units and maintain the weight magnitudes used in Jercog et al. (2017), we normalized the observed *θ* and *g* to those of the excitatory unit used by Jercog et al. (*θ_E_*=5, *g_E_*=1), leading to *θ_X_* values of 5, 30, 15, and *g_x_* values of 1,2.7, and 1.6 for the E, P, and S, populations respectively.

We also set the values of *τ_x_* to the fits of the mean membrane time constants of the Pyr, PV, and SST neurons (10, 4, and 6 ms for *E, P*, and *S*, respectively). We note, however, that the time constant in Eq. 1–3 is often interpreted as corresponding to the synaptic time constant (Shriki et al., 2003; Ozeki et al., 2009), rather than the membrane time constants. However, this interpretation is less parsimonious with the current formulation in which the IPSCs from PV and SST cells can have different time constants.

### Fits of the model weights to the empirically-defined firing rates

In order to find the weights sets or vectors that captured the empirically observed mean firing rates of the Pyr, PV, and SST neurons, we performed a parameter search over the 9 weight variables with empirically-derived values of, i.e., *θ_E_, θ_P_, θ_S_, g_E_, g_P_, g_S_, τ τ_P_*, and *τ_S_*. The search values were approximately centered on the values used by Jercog et al (2017):

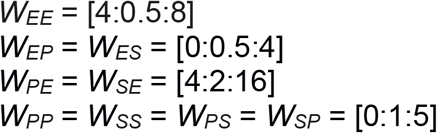

The bracketed values represent the minimum, step size, and maximal values of the corresponding weight search. Overall, we searched a parameter space of 9×9×9×7×6×6×7×6×6 for a total of 46,294,416 parameter sets. We defined the accepted group of weight sets as those which led to fits in which all three Up-state firing rates (i.e., steady-state E, P, and *S* values) where within ±25% of the empirically observed rates (5, 14, and 17 Hz, for the Pyr, PV, and SST populations, respectively).

### Analysis of fixed-points and the paradoxical effect

In order to derive the Up-state fixed points for all three populations (*E*, P*\*, and *S*\*) of Eqs. 1-3 in the linear regime (i.e., *E, P*, and *S* > 0), we can first impose that all three derivatives be equal to 0 and obtain:

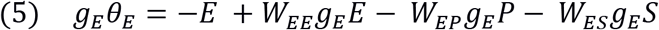

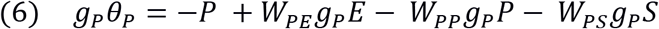

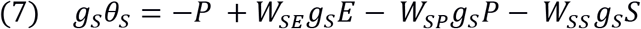

By solving this set of equations, and through multiple substitution steps, it can be shown that the Up-state fixed point of *P* is:

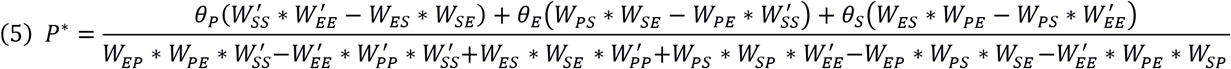

where 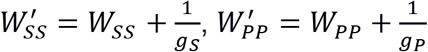, and 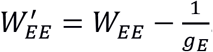. Importantly, it is only the first term of the numerator that is critical to determining whether the paradoxical effect (in the *P* population) is present in response to *P* activation. Specifically, external tonic *depolarization* of the *P* population is equivalent to *decreasing θ_P_*. Thus we can see that such a decrease will lead to a decrease in *P*\* (i.e., the paradoxical effect) if 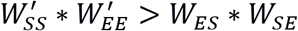, and an increase in *P\** if 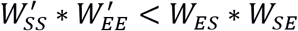. An equivalent equation can be derived for *S*\*.

## RESULTS

### The two-population model of Up-states

Up-states are characterized by transitions from a quiescent network-wide Down-state to a regime in which excitatory neurons are depolarized and fire at moderate rates. **Fig. 1A** illustrates Up-Down transitions in two simultaneously recorded pyramidal neurons (Pyr) in an *ex vivo* slice of the auditory cortex. Both neurons transition between Up- and Down-states more or less simultaneously, and during Up-states maintain an approximately constant level of membrane depolarization resulting in the characteristic bimodal distribution of membrane voltages (Mann et al., 2009; Lőrincz et al., 2015). Importantly, during Up-states, the firing rate of pyramidal neurons is far below saturation, indicating that they do not reflect pathological regimes characterized by saturated firing rates or paroxysmal discharges (Connors, 1984; Prince and Tseng, 1993; Timofeev et al., 2004).

**Figure 1.**
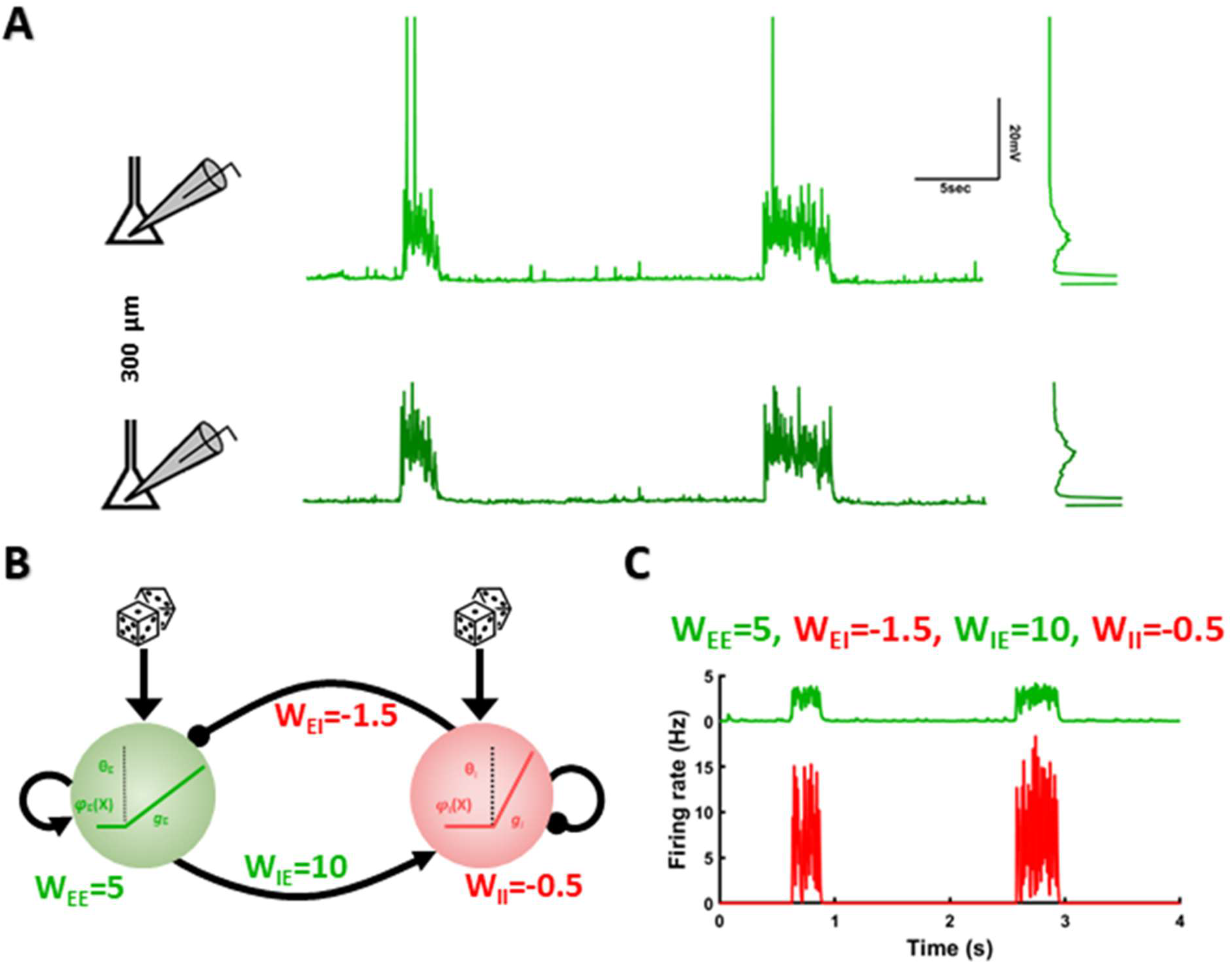
Experimental Up-states and numerical simulations in a two-population model. **(A)** Example of Up states recorded in an *ex vivo* cortical circuit. Traces represent two simultaneously recorded pyramidal neurons separated by approximately 300 μm. Insets to the right show the bimodal distribution of membrane voltage (the peak of the distribution which is centered at rest, is clipped for visualization purposes). **(B)** Schematic of the two-population model with an excitatory (green) and inhibitory (red) population. Each population is simulated by a linear threshold activation function defined by threshold (*θ*), and gain (*g*). Lines with arrows represent excitatory connections and lines with dots represent inhibitory connections. Dice represents noise. **(C)** Example of simulated Up-states in the E (upper trace) and I populations (bottom trace) from the model with the indicated weight values.

The Up- and Down-state transitions illustrated in **Fig. 1A**, have typically been interpreted to reflect bi-stable network regimes in which Down-states correspond to a trivial fixed-point where excitatory firing rates are approximately zero, and Up-states correspond to a nontrivial fixed-point with moderate firing rates. Computational studies have carefully characterized such regimes in firing rate and spiking neural networks models (Brunel, 2000; Holcman and Tsodyks, 2006; Renart et al., 2010; Dao Duc et al., 2015; Jercog et al., 2017; Tartaglia and Brunel, 2017). **Fig. 1B** shows an example of a firing rate model of Up-states based on Jercog et al. (2017; see Methods). The model is composed of variables representing the firing rates of a population of excitatory (E) and inhibitory *(I)* neurons connected through the synaptic weights: *W_EE_* (*E→E*), *W_EI_* (*I→E*), *W_IE_* (*E→I*), *W_II_* (*I→I*). Intuitively, the Up- and Down-state transitions in the model can be understood as a noisy excitatory input to the *E* population, which sometimes reaches threshold, and triggers positive feedback through the *E→E* connection. In parallel, the *I* population receives an increasing excitatory drive from *E*, triggering rapid and strong inhibition of the *E* unit—settling in an inhibition stabilized fixed-point attractor (**Fig. 1C**) (Tsodyks et al., 1997; Ozeki et al., 2009; Jercog et al., 2017).

As with most Up-states models to date, this model incorporates only a single unspecified class of inhibitory neurons. However, it is well established that there are distinct populations of inhibitory neurons within the neocortex and that these neurons have distinct properties and functional roles in cortical computations (Adesnik et al., 2012; Kuhlman et al., 2013; Pi et al., 2013; Natan et al., 2017). Here we focused explicitly on PV and SST neurons as they are the most common types of inhibitory neurons (Rudy et al., 2011), and because VIP neurons have been associated with changes in brain states and cross-cortical interactions (Pi et al., 2013; Krabbe et al., 2019), which are not present in the *ex vivo* experimental preparation we are simulating.

### Characterization of Pyr, PV, and SST neurons during Up-states

To characterize the firing rates and intrinsic properties of PV and SST neurons, we recorded from *ex vivo* cortical slices of PV-Cre and SST-Cre animals that expressed the tdTomato marker in PV or SST neurons, respectively. Targeted paired whole-cell recordings of Pyr and PV, or Pyr and SST were performed. Both PV and SST neurons transitioned between Up- and Down-states in synchrony with Pyr neurons (**Fig. 2A-B**). PV and SST neurons exhibited firing rates significantly above those of the Pyr neurons (**Fig. 2C**, one-way ANOVA, F_2,75_ = 20.3, p<0.001). The median firing rate of the three neuron classes was: Pyr: 4.5, IQR=6.09 Hz (n=39), PV: 14, IQR=8.55 Hz (n=26) and SST: 17, IQR=14.58 Hz (n=13) (Pyr x PV: Wilcoxon signed-rank, z = – 5.02, p < 0.001; Pyr x SST, z = −4.19, p < 0.001).

**Figure 2.**
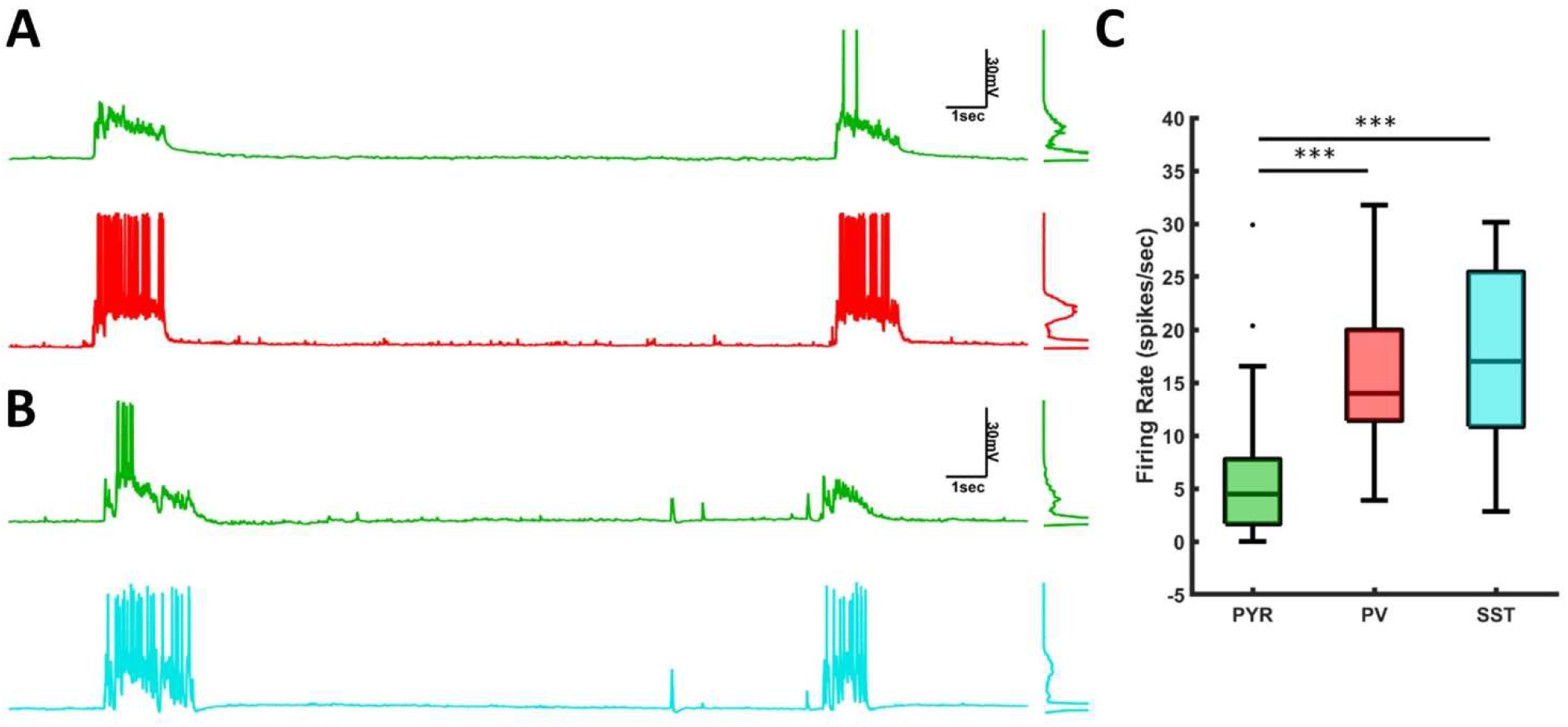
Experimentally-observed Up-state signatures in Pyr, PV, and SST neurons. **(A)** Example of Up-states in simultaneously recorded Pyr and PV neurons. **(B)** Example of Up-states in simultaneously recorded Pyr and SST neurons. **(C)** Mean firing rate of Pyr, PV, and SST neurons during Up-states.

In order to build a three-population model composed of an excitatory population and two inhibitory neuron populations, a critical issue is what are defining the differential characteristics of the two inhibitory neuron populations. Thus, as a first step towards building an empirically grounded three-population model of Up-states, we measured the Frequency-Current (F-I) function of Pyr, PV, and SST neurons and characterized their firing properties (**Fig. 3A**). We fit the F-I curves to a rectified linear unit (ReLU) function, defined by two variables: a threshold (*θ*) and slope (the gain g). The fitted threshold (*θ_E_*) and gain (gE) values of the Pyr neurons (*θ_E_*=0.06 nA, *g_E_*=124 Hz/nA) were below the threshold and gain values of the PV (*θ_P_*=0.36 nA, *g_P_*=334 Hz/nA) and SST (*θ_S_*=0.18 nA, *g_S_*=198 Hz/nA) neurons (**Fig. 3B**).

**Figure 3.**
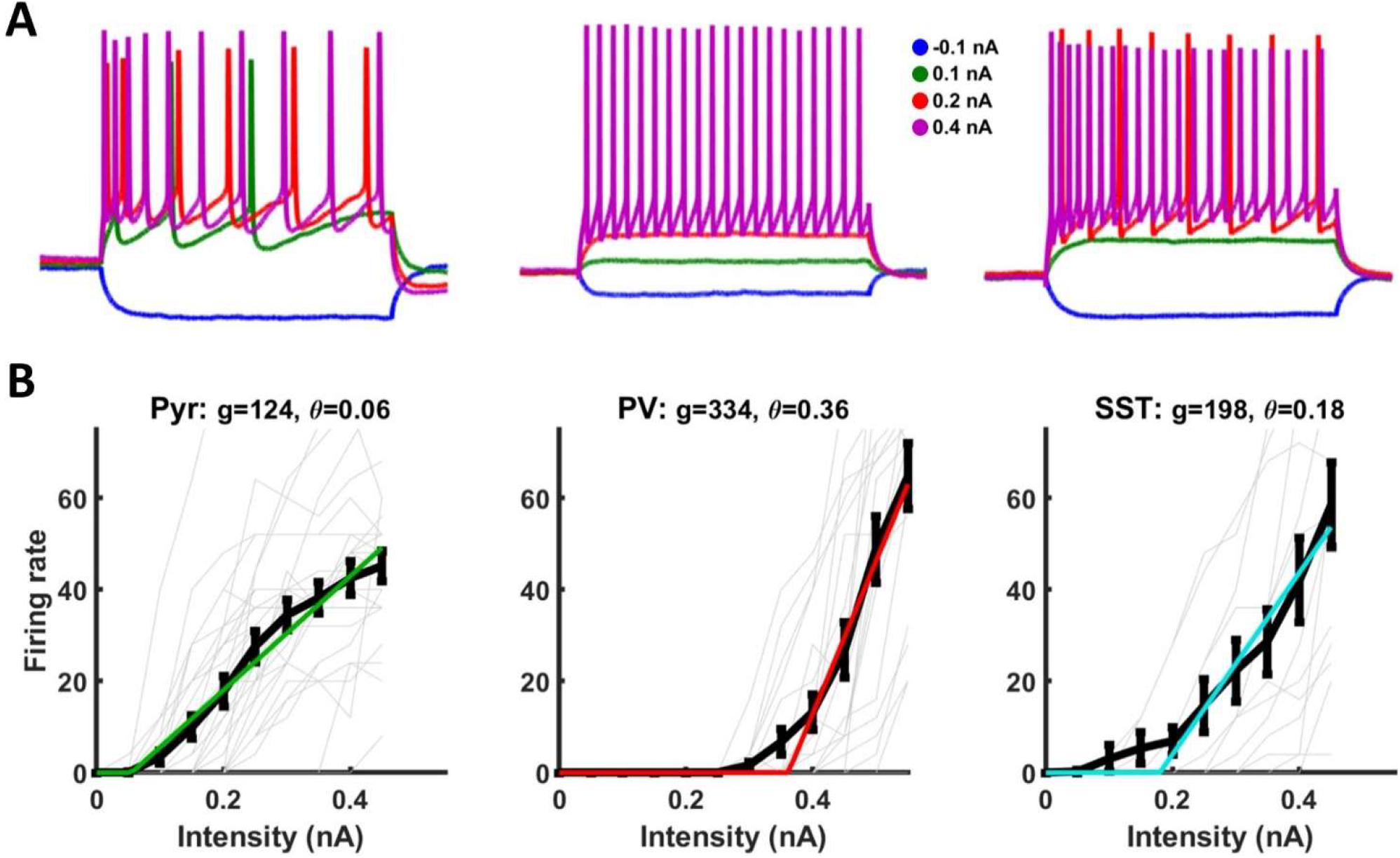
F-I curve fits of the intrinsic excitability of Pyr, PV, and SST neurons. **(A)** Top: Sample intrinsic excitability of Pyr (left), PV(middle), and SST(right) neurons in response to depolarizing current steps of −0.1,0.1,0.2, 0.4 nA. **(B)** Threshold-linear fits of the F-I curves of Pyr populations. Light gray lines are the F-I curves of individual neurons. Black lines are the mean plus standard errors of the F-I curves, and colored lines are the fits. The gain (*g_P_*=334) and the threshold (*θ_P_*=0.36) of PV (middle) neurons is higher than in Pyr (*g_E_*=124, *θ_E_*=0.06; left) and SST neurons (*g_S_*=198, *θ_S_*=0.18; right).

### Empirically-based Up-state model with Pyr, PV, and SST populations

To the best of our knowledge, the above results provide the first opportunity to directly simulate a three-population model based on empirically defined F-I curves and fit the weights of the model to match the observed firing rates in the three neuronal populations. We thus extended the model presented in **Fig. 1B**, to include three populations: Pyr (*E*), PV (P), and SST (S) (**Fig. 4A**). While *in vitro* experiments from acute slices have revealed a significant amount of information about the interconnectivity between these three populations (Silberberg and Markram, 2007; Pfeffer et al., 2013; Xu et al., 2013), we chose to perform an unbiased assumption-free implementation of the model to directly determine the predictions generated by a model based on empirically-derived F-I curves. Thus, we implemented a fully interconnected model with all nine connection classes (**Fig. 4A**, see Methods).

**Figure 4.**
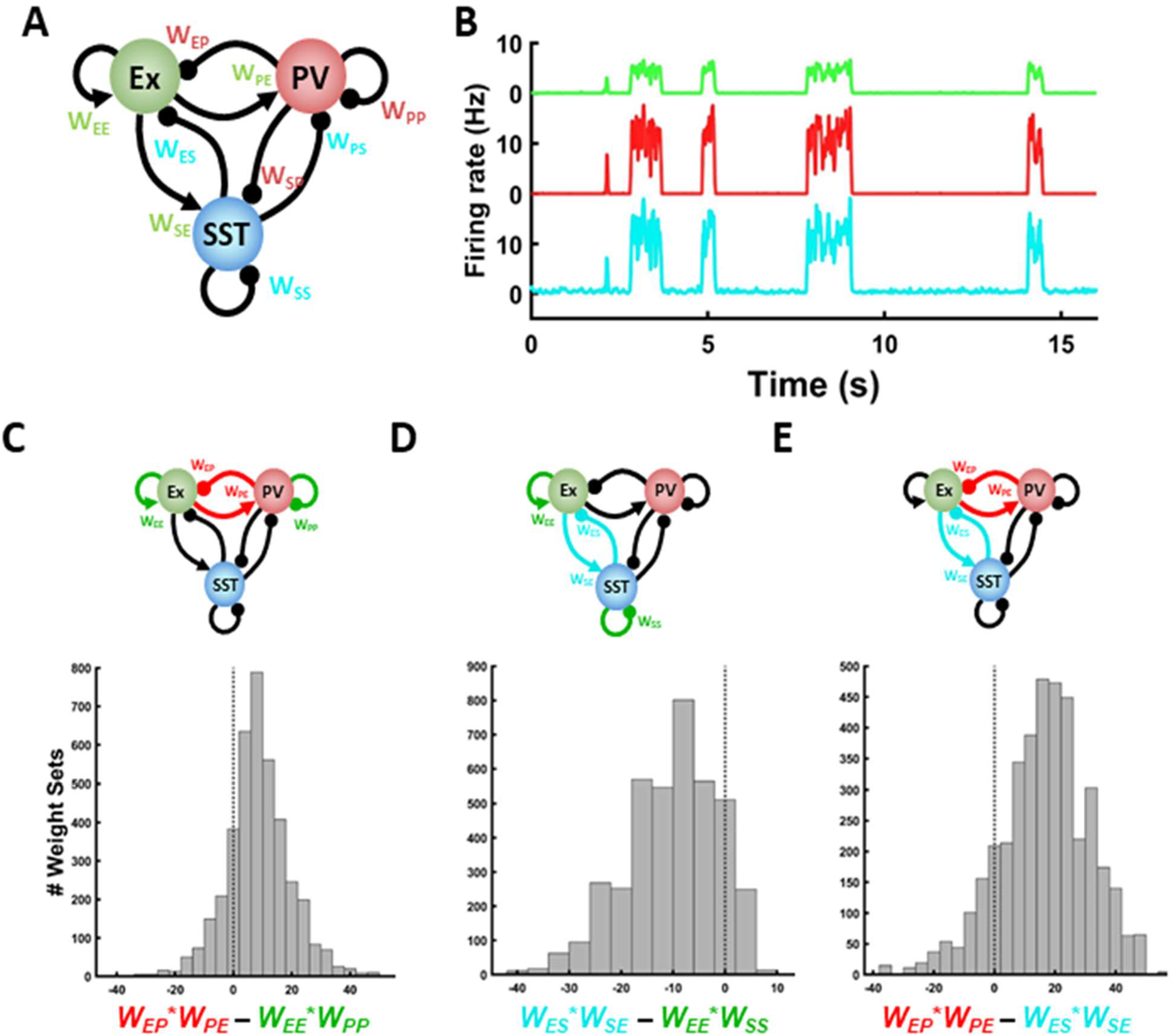
Empirically-derived three-population model of Up-states. **(A)** Schematic of the fully connected three-population model. Figure conventions are similar to **Figure 1B**. **(B)** Example of Down↔Up state transitions in the model of Pyr (upper), PV (middle) and SST (bottom) units. The weights were chosen as the set closest to the centroid of the distribution of the fit weight sets. **(C)** Distribution of the *fit weight set* (i.e., those the weight vectors that captured the experimentally observed firing rates) of the inhibitory (red) minus the excitatory (green) loop of the Pyr and PV interaction. **(D)** Distribution of the weights of the inhibitory (cyan) minus the excitatory (green) loop of the Pyr and SST interaction. **(E)** Distribution of the weights of the inhibitory (red) loop of the Pyr-PV minus the inhibitory (cyan) loop of the Pyr-SST interaction.

We performed an unbiased search to determine which sets of weights would generate the experimentally observed firing rates (see Methods). By an unbiased search, we mean that the parameter space was the same for all *P* and *S* related weights—e.g., the range of *W_EP_* values explored was equal to the range of *W_ES_* values. We explored approximately 46 million sets of weights. As there are nine weights and three firing rates to be fit, the parameter search is underconstrained, resulting in a distribution of weight sets that accounted for the experimentally observed firing rates. Out of 46 million weight sets, approximately 4,000 generated Up-state firing rates of all three cell types (i.e., steady-state *E, P*, and *S* values) that were within ±25% of the empirically observed rates. We refer to this as the set of *fit weights* (see Methods). **Fig. 4B** shows spontaneous Up- and Down-states of the E, P, and *S* populations in a simulation run with the prototypical weight set—which we define as the weight vector closest to the centroid of the all fit weights vectors.

To test the hypothesis that the empirically-derived intrinsic properties of the PV and SST cells play a deterministic role in their functional contribution to Up-states, we explored the relative distribution of weights of the *P* and *S* populations in the fit weight sets. Numerical and analytical analyses have established that in the two-population model a necessary condition for the existence of Up-states is that the net inhibition be stronger than net excitation. Mathematically this condition is expressed as *W_EI_W_IE_* > *W_EE_W_II_* (ignoring the contribution of the gains). Note that *W_EI_W_IE_* reflects the strength of the net inhibitory loop in the two population model, while *W_EE_W_II_* reflects net excitation as the weight of the inhibitory population onto itself (*W_II_*) is functionally excitatory. We thus first compared the distribution of the net inhibition minus the net excitation for the *E↔P* and *E↔S* loops. As shown in **Fig. 4C-D** in 80% of the fit weight sets, the *E↔P* loop satisfied the criteria *W_EP_W_PE_* > *W_EE_W_PP_*. In contrast, in the *E↔S* loop only 8% of the fit weights was *W_E_ W_SE_>W_EE_W_SS_*. To directly contrast the contribution of *P* and *S* populations we also examined the net *P* inhibition minus the net *S* inhibition: *W_EP_W_PE_ – W_ES_W_SE_*. This contrast revealed that in 85% of the cases, the net *P* inhibition was more robust than the net *S* inhibition (**Fig 4E**). These results indicate that solely due to the intrinsic properties of the two inhibitory neuron classes, the *P* population generally emerges as the one primarily responsible for an inhibition stabilization. Nevertheless, in a minority of the parameter regimes, *W_ES_W_SE_* was larger than *W_EP_W_PE_*; however, in most of these cases, *W_EP_* or *W_PE_* was close to or equal to zero. Thus in the absence of a *P↔E* loop, the intrinsic properties of the *S* neurons form a “backup” system that can support Up-states and fulfill the role of inhibition stabilization.

### The Paradoxical Effect

The above results predict that the net inhibition in the *E↔P* loop is stronger than in the E↔S loop. But interestingly, this observation does not translate into meaning that PV neurons are more effective at modulating Up-states—e.g., turning Up-states off. In part precisely because of the strength of the *E↔P* loop, changes in the external input to *P* engage a dynamic rebalancing of excitation and inhibition resulting in counterintuitive properties, including the so-called *paradoxical effect* that has been described in the two-population model (Tsodyks et al., 1997; Rubin et al., 2015). Specifically, in the two-population model (e.g. **Fig. 1B**) if the *I* unit is artificially activated during an Up-state, an overall *decrease* in *I* activity is observed. This is because increasing *I* decreases E, which in turn decreases excitatory drive to *I.* The net result is that for *E* to settle at a lower fixed point, *I* must also settle at a lower fixed point (to maintain an approximate E/I balance). In the two-population model, this paradoxical effect is obligatory (Tsodyks et al., 1997; Jercog et al., 2017). Thus, we next examined the properties of the paradoxical effect in the three-population model and whether *P* or *S* activation is more effective at turning off Up-states (**Fig 5A**). In a model with the prototypical weights, weak *P* activation during an Up-state (see Methods) produced a paradoxical effect. In contrast, weak *S* stimulation did not result in a paradoxical effect, i.e., there was an increase in steady-state *S* activity after external *S* activation (**Fig 5A, middle panel**). Across the 3963 weight sets that fit the data, the paradoxical effect was observed in 95% and 18% of cases in the *P* and *S* units, respectively (**Fig. 5A, bottom panel**).

**Figure 5.**
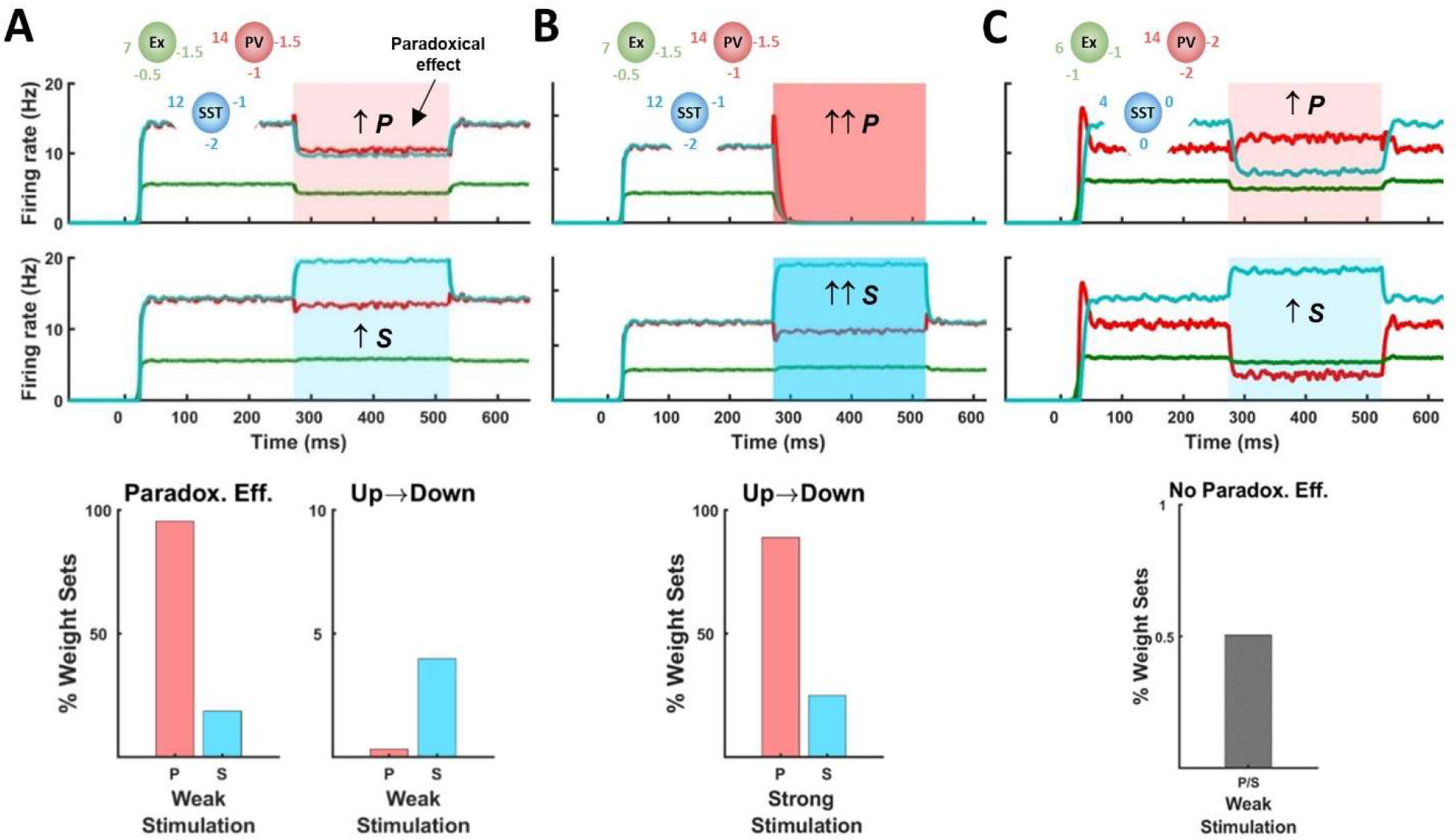
The paradoxical effect in the three-population model. **(A) Top**: Example of weak excitation of the *P* units during an evoked Up-state (upper panel; red overlay) showing the paradoxical effect—i.e., despite depolarizing the *P* unit a decrease in *P* firing rate is observed. In contrast, weak depolarization of the *S* unit (lower panel; cyan overlay) did not produce a paradoxical effect in the *S* unit. **Bottom:** In response to weak activation of *P* units the paradoxical effect was observed in the *P* units across the vast majority of fit weight sets (left bar of left panel). In contrast, the paradoxical effects were generally not observed in the *S* units in response to weak *S* activation (right bar of left panel). However, *S* activation was more likely to produce Up→Down transitions (right panel). **(B)** Top: Same as **C** with strong activation of *P* (upper) and *S* (lower). Bottom: Strong activation of *P* cells was much more effective at inducing Up→Down transitions across of the validated weights sets. **(C) Top**: Example of a parameter regime in which the paradoxical effect is not observed in either *P* (upper) or *S* (lower) units. Note that in this regime small increases in *P* (*S*) results in relatively larger decreases in *S* (*P*). **Bottom**: Regimes in which the paradoxical effect is not observed in both *P* and *S* units comprise a very small subset of the total fit weight set.

Counterintuitively, although the *E↔P* inhibitory loop was stronger than the E↔S loop, weak activation of *S* units was actually more likely to induce an Up→Down transition than weak activation of *P* units (**Fig. 5A, bottom panel**). But as expected, strong depolarization of *P* or *S* (**Fig. 5B**) units was likely to induce Up→Down transitions, and now *P* activation was more effective at turning off Up-states (89%) than *S* activation (25%, **Fig. 5B, bottom panel**).

In the vast majority of the fit weight vectors, the paradoxical effect was observed in either the *P* or *S* units in response to *P* or *S* stimulation, respectively. Interestingly, however, a very small number of weight sets (0.5%) in which the paradoxical effect was not observed in either the *P* or *S* population (**Fig. 5C**). A fairly strong PV↔SST interaction characterized these regimes. This finding has an important experimental implication as it establishes that although the paradoxical effect is expected in most parameter regimes, it is not obligatory in a model with two types of inhibitory neurons—a potentially relevant observation because a number of experimental studies have failed to observe the paradoxical effect (Xu et al., 2013; Gutnisky et al., 2017, see Discussion).

Analysis of the steady-state equations confirmed the numerical simulations (see Methods), and showed that whether or not the paradoxical effect is present in the *P* population, for example, depends on the coupling between the *E* and *S* populations. Specifically, if 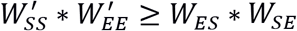 (i.e., if excitation is stronger than inhibition in the *E↔S* loop, see Methods) the paradoxical effect will be observed in the *P* population. Indeed, we can see that if *W_ES_* or *W_SE_* is equal to zero, the network is effectively equivalent to the two-population model, in which case the paradoxical effect must be present in the *P* population. In contrast, if *W_ES_* * *W_SE_* is relatively high (strong *S* inhibition), then the *S* population will be responsible for network stabilization and the paradoxical effect will not be observed in the *P* population.

Overall, these simulations lead to a number of experimental predictions, including: 1) that the Pyr↔PV loop should be stronger than the Pyr↔SST loop; and 2) that strong PV activation should be more effective than SST activation at turning off Up-states. Below we address these two predictions using *ex vivo* cortical circuits.

### Testing predictions of the model: paired Pyr-PV and Pyr-SST recordings

To directly examine the predictions of the model, we first performed paired recordings from Pyr-PV and Pyr-SST neuron pairs using *ex vivo* cultures from PV-Cre and PV-SST mice, respectively. As shown in **Fig. 6,** the observed connection probability was consistent with the predictions of the model—note that because the model is population-based, the weights reflect both connection probabilities and mean synaptic strengths. Of 53 Pyr-PV and 27 Pyr-SST pairs, we found connections in approximately 50% and 25% of them, respectively. The Pyr→PV connection probability was 0.31, while the PV→Pyr was 0.23 (**Fig. 6B**). Interestingly, reciprocal connections were more likely than expected from the independent probabilities, indicating that reciprocity is a favored motif (Song et al., 2005). Most of the detected connections involving SST neurons were in the Pyr→SST direction (connection probability = 0.23) (**Fig. 6C**). However, we stress that the low probability (0.04) of SST→Pyr connections is likely an underestimation because SST synapses are generally on dendrites and that we performed the experiments in current-clamp—making it difficult to detect weak inhibitory connections. The average unitary Pyr→PV amplitude (2.87±0.82 mV) was significantly larger than the unitary Pyr→SST strength (0.70±0.21 mV). The mean PV→Pyr amplitude was (1.93±0.47 mV) larger than the single connected SST→Pyr unitary IPSP amplitude we recorded (0.3 mV). Although these results are limited in their ability to provide quantitative estimates for the model parameters, they confirm the model prediction that the Pyr↔PV inhibitory loop is stronger than the Pyr↔SST loop in the *ex vivo* system we simulated.

**Figure 6.**
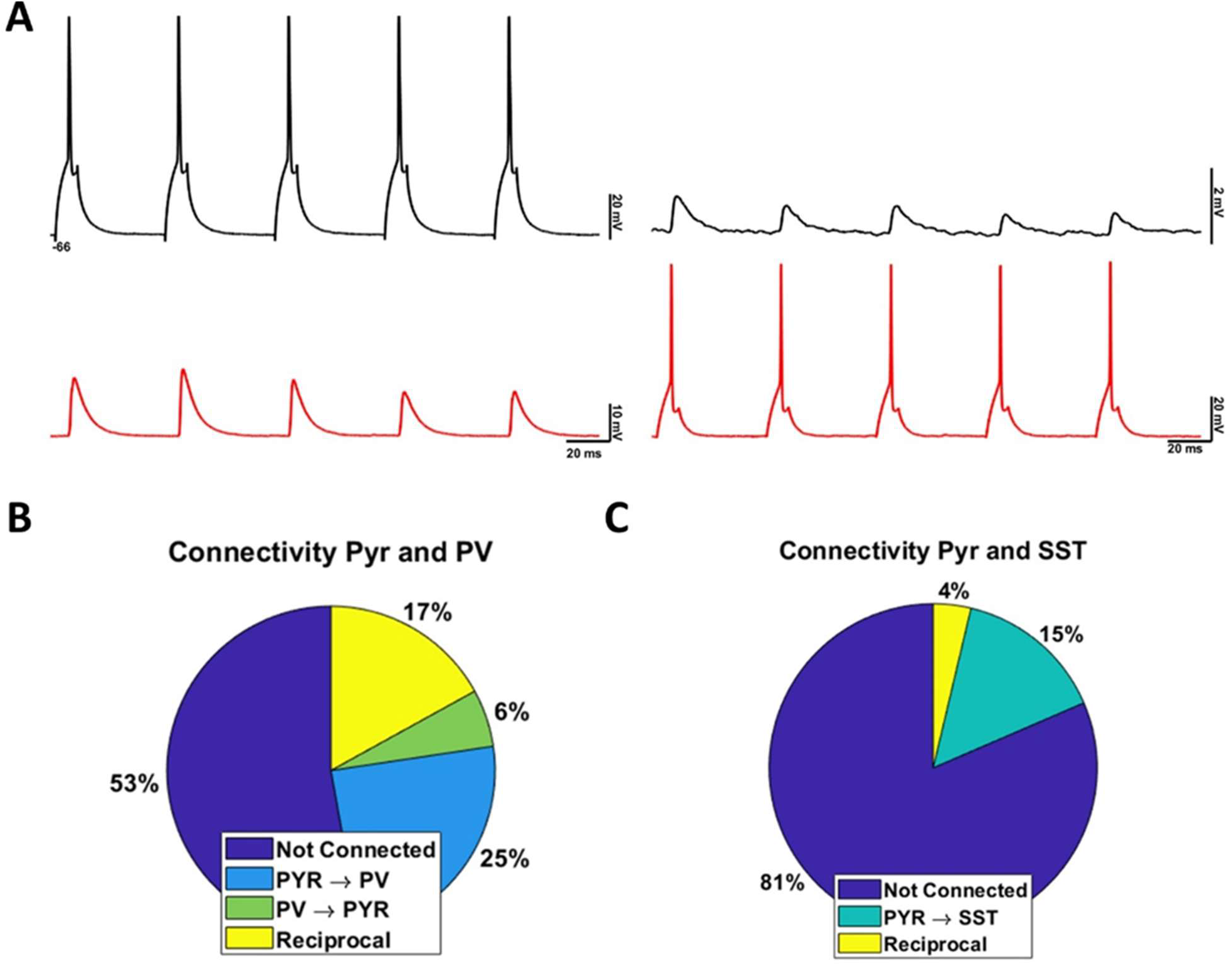
Paired recordings from Pyr-PV and Pyr-SST neuron pairs. **(A**) Simultaneously recording of Pyr and PV neurons. Spikes in the Pyr neuron (upper left trace) induce EPSPs in the PV interneuron (bottom left trace) while spikes in PV interneuron (bottom right trace) induced IPSPs in the Pyr neuron (upper right trace). **(B)** Pie chart of connectivity of Pyr-PV pairs. **(C)** Same as B but for Pyr-SST neuron pairs.

### Testing predictions of the model: activation of PV but not SST units turns off Up-states

As shown in **Fig. 5B,** the model predicts that strong activation of PV neurons during Up-states should be more effective at turning off Up-states than activation of SST neurons. To test this prediction, we expressed ChETA in PV or SST cells in separate *ex vivo* slices and examined whether optogenetic activation of these inhibitory neuron populations produced Up→Down transitions. Towards this end, we recorded from Pyr neurons and triggered a train of light pulses during detected Up-state onsets in slices expressing ChETA in either PV or SST neurons (**Fig. 7**; see Methods). We used “full-field” light activation to emulate the strong activation of the model (**Fig. 5B**) and to be able to further increase the already high firing rate of inhibitory neurons during Up-states (**Fig. 2**). Optogenetic stimulation in PV and SST neurons effectively increased inhibitory neuron firing rate even during Up-states (**Extended Fig. 7-1**). The comparison of light-off and light-on trials revealed that activation of PV neurons during Up-states was highly effective at inducing Up→Down transitions (mean Up state duration: light off, 1.80 ± 0.36 sec, light on, 0.78 ± 0.34 sec; n=10, Wilcoxon signed-rank test: z = 2.80, p = 0.005) (**Fig. 7B**). Similarly, optical activation of PV neurons also produced significant decrease in the voltage of the pyramidal neuron (−56.26 ± 1.93 mV versus −64.12 ± 1.64 mV; z = 2.80, p=0.005) (**Extended Fig. 7-2**). In contrast, SST activation did not produce a significant decrease in Up-state duration (1.76 ± 0.19 sec, n=21; 2.12 ± 0.22 sec, n=21, z-stat = −1.09, p = 0.28) (**Fig. 7D**), nor in the voltage of the pyramidal neuron (**Extended Fig. 7-2**). These results further confirm the prediction of the model, that because Pyr-PV connectivity is stronger than Pyr-SST connectivity, strong activation of PV neurons is more effective at inducing Up→Down transitions. Indeed, these experimental results are in agreement with the prototypical weight set simulations shown in **Fig. 5B**, in which strong *P* activation produced an Up→Down transition, and *S* activation had minimal effect on *E* activity.

**Figure 7.**
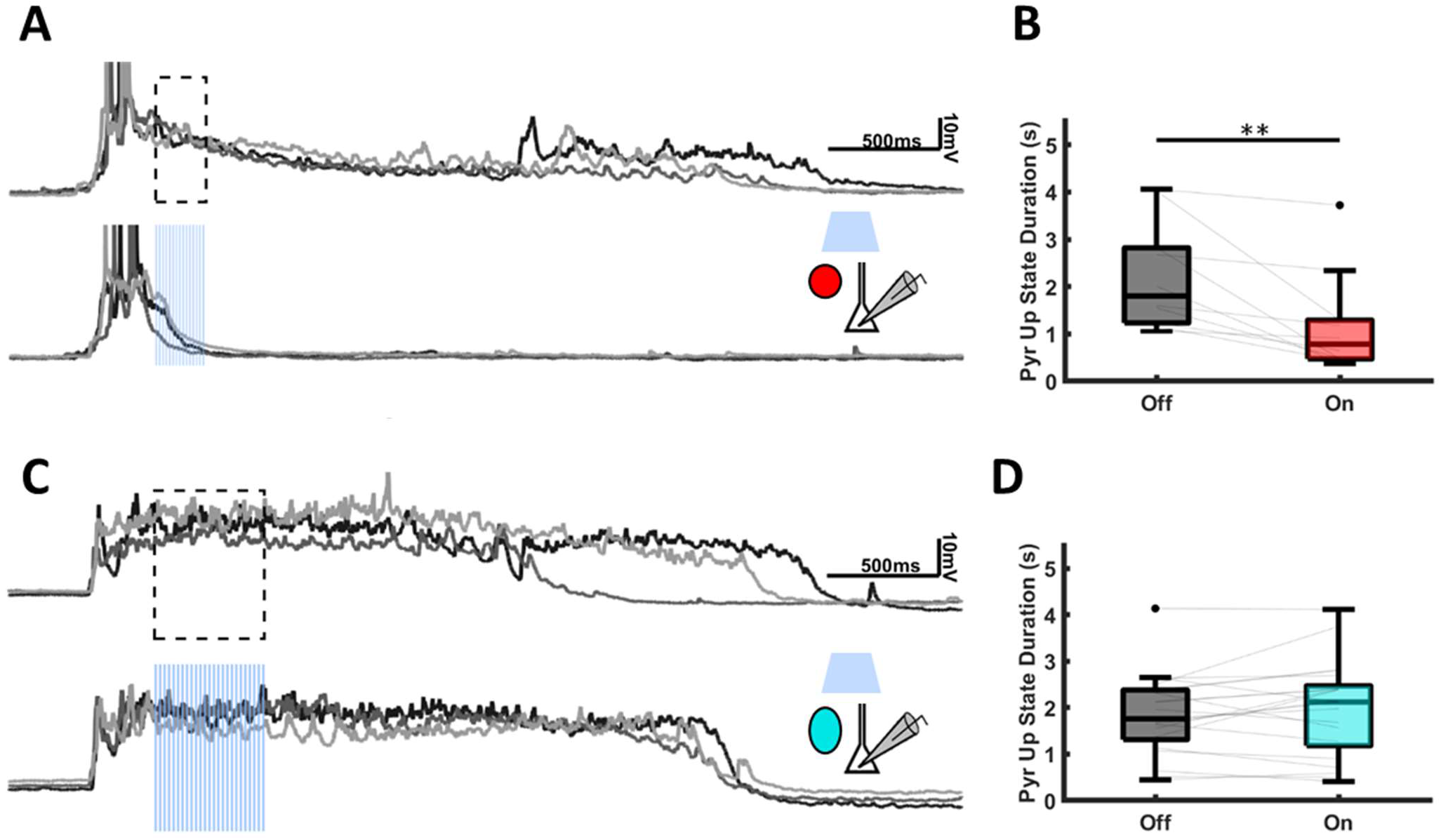
Differential effects of PV and SST interneurons to turn off Up-states. **(A)** PV light activation resulted in shorter Up states as depicted by individual traces (bottom gray) and the mean Up-state trace (bottom black) compared to Up-states in which light was not presented (Upper traces). **(B)** Median Pyr Up-state duration was significantly shorter when the light was delivered during Up-states compared to Up-states that did not receive light stimulation **(C)** Same as A but for SST neurons. Up-states that were presented with light are not different than Unstimulated Up-states. **(D)** SST activation during Up-states resulted in similar Up-states duration compared to unstimulated Up-states.

## DISCUSSION

We experimentally characterized the intrinsic input-output function (the F-I curve) of three classes of neurons in *ex vivo* organotypic cortical cultures. We incorporated these empirically-derived activation functions into a computational model that captures the experimentally observed firing rates, and finally, tested experimental predictions of the model. This integrated experimental-computational approach parallels studies in invertebrate systems (e.g., Gingrich and Byrne, 1985; Buonomano et al., 1990; Marder and Calabrese, 1996; Phares et al., 2003; Prinz et al., 2004). Such approaches, however, have been difficult to carry out in neocortical circuits for a number of reasons, including that the network regimes being simulated are often recorded *in vivo* while the cellular properties are generally obtained from *in vitro* acute slice preparations (and thus under significantly different physiological conditions and network regimes). And in contrast to acute slice studies in which the state of neurons and synapses reflects the homeostatic learning rules shaped by the *in vivo* environment, here we studied *ex vivo* circuits that converged to their homeostatic set-points over weeks. Furthermore, all *in vivo* data sets are subject to reciprocal influences from a multitude of brain areas that cannot be incorporated into models. Thus, by studying and modeling an *ex vivo* cortical circuit, it was possible to directly couple the experimental data to the computational model and ensure that the experimental and computational results reflect the properties of local intrinsic circuits.

### Differential role of PV and SST neurons in network stabilization

Our findings establish that the empirically-derived differences in F-I characteristics of PV and SST neurons are sufficient to lead to a pronounced differential contribution of these neuron classes to network dynamics. The threshold and gain of PV neurons were higher than that of SST neurons. This signature is critical to the stabilization of Up-states for two reasons: 1) the increased threshold means that the excitatory neurons can engage in positive feedback at very low firing-rates before triggering inhibition; 2) once the inhibitory neurons reach threshold they can quickly overtake the activity in the excitatory neurons because of their larger gain (Jercog et al., 2017), thus stabilizing the network. Interestingly, while the differential gain between PV and SST neurons has not been carefully contrasted in the past, the threshold difference in PV and SST neurons is a defining distinction between them, e.g, so-called low-threshold inhibitory neurons correspond to SST neurons (Gupta et al., 2000; Rudy et al., 2011; Cardin, 2018). Thus, our results emphasize that a critical functional difference between PV and SST neurons is that both the threshold and gain of the F-I function of PV neurons are larger than those of SST neurons.

It is generally accepted that the observed diversity of inhibitory neurons reflects that cortical computations require inhibition to fulfil multiple functional roles (Wang et al., 2004; Pfeffer et al., 2013; Natan et al., 2015). However, while the differential contribution of these neurons has focused primarily on their synaptic and connectivity properties, our results suggest a critical role of the differential input-output function of inhibitory neuron subclasses in cortical dynamics. Furthermore, the importance of the difference in the intrinsic excitability properties of PV and SST neurons is consistent with the importance of intrinsic excitability in neural computations (Marder and Calabrese, 1996; Josselyn and Tonegawa, 2020). However, an open question pertains to the degree to which the input-output functions of PV and SST neurons are plastic. While it seems likely that the threshold and gain parameters of these neural populations undergoes some forms of intrinsic plasticity (Daoudal and Debanne, 2003; Zhang and Linden, 2003; Debanne et al., 2019), we would hypothesize that fundamental properties reported here— i.e., *θ_P_ > θ_S_* and *g_P_* > *g_S_*—reflect fundamental and hard-wired differences within these populations.

Because the experimental data was collected in *ex vivo* cortical circuits our results provide insights into the unsupervised learning rules governing the emergence of Up-states. Specifically, Up-states seem to reflect states that cortical circuits actively seek out through homeostatic mechanisms. For example, while cortical cultures are initially silent, spontaneous and evoked Up-states emerge over the course of *ex vivo* development (van Pelt et al., 2005; Johnson and Buonomano, 2007; Motanis and Buonomano, 2015; Motanis and Buonomano, 2020). This suggests a set of learning rules in place *ex vivo* that drive networks towards Up-Down state transitions similar to those observed *in vivo* and in acute slices (Sanchez-Vives and McCormick, 2000). Given the critical stabilizing effect of PV neurons, our results suggest that these learning rules may operate primarily on PV neurons.

The computational model predicted that the strength of the Pyr↔PV inhibitory loop is significantly stronger than the Pyr*↔*SST loop. Paired recordings and optogenetic experiments validated these predictions. These results, however, do not imply that *in vivo* SST neurons do not play a stronger role in Up-state dynamics. An *in vivo* study, for example, reported that both PV and SST activation could terminate Up-states (Zucca et al., 2017). Similarly, studies in acute slices have also directly implicated SST neurons in Up-state dynamics (Urban-Ciecko et al., 2015; Neske and Connors, 2016). Indeed, given the ability of cortical circuits to adapt to a wide range of natural and pathological regimes, it is expected that different inhibitory neurons play different roles in different contexts. However, our results reveal that under highly controlled conditions and in the absence of external input from other brain areas, cortical microcircuits converge to regimes where PV neurons are primarily responsible for inhibition stabilization emergent dynamic regimes. Thus, we would suggest that future research on the learning rules governing network dynamics should focus primarily on the homeostatic plasticity of the Pyr ↔PV loop.

### The Paradoxical Effect and Predictions

The standard two-population model of Up-states predicts the presence of a paradoxical effect, in which external depolarization of the inhibitory neuron population during an Up-state will counterintuitively decrease the firing rate of inhibitory neurons. Testing this prediction has been experimentally challenging for a number of reasons, including the need to record and carefully control the levels of external input to a specific inhibitory neuron population during Up-states (Sanzeni et al., 2020). Some experiments have reported signatures consistent with the paradoxical effect (Kato et al., 2017; Moore et al., 2018; Sanzeni et al., 2020), while others have not (Xu et al., 2013; Gutnisky et al., 2017). However, until recently, computational models have not explicitly addressed whether the paradoxical effect is present in the context of a three-population model that included two inhibitory neuron classes. A recent model of asynchronous cortical activity has shown that a paradoxical effect is not obligatory in the PV population in a model that included both PV and SST neurons (Sanzeni et al., 2020). Our analysis of the three-population model of Up-states shows that whether the paradoxical effect is present or not in PV neurons depends on the relationship of the excitatory (*W_EE_*W_SS_*) and inhibitory loop (*W_ES_*W_SE_*) between the Pyr and SST population. Our numerical results reveal that in the vast majority of cases the paradoxical effect was observed in either the PV or SST population (**Fig. 5**), but there is a very narrow regime in which neither population of inhibitory neurons revealed a paradoxical effect. Based on our numerical simulations we predict that the paradoxical effect should be observed in PV neurons, although it is possible that some failures to observe the paradoxical effect in PV neurons could reflect situations in which the SST are driving network stabilization, in which case they should exhibit the paradoxical effect. One must also consider the possibility that experimental failures to observe the paradoxical effect may be because the standard computational models of Up-states, including ours, lack critical biological ingredients. For example, most models do not explicitly distinguish between the contribution of GABA_A_ and GABA_B_ receptor-mediated inhibition, yet GABAB-mediated inhibition powerfully regulates Upstates (Mann et al., 2009; Urban-Ciecko et al., 2015). Future work will attempt to explicitly address the paradoxical effect in both PV and SST neurons in our preparation and the potential contribution of other mechanisms to Up-state dynamics.

## Conclusions

Using an *ex vivo* preparation, it was possible to use empirically derived fits of the F-I functions of PV and SST neurons to create a model of Up-state dynamics and fit the weights of the model to the empirically derived Up-state firing rates. This approach allowed for a direct link between the experimental and computational components. The results establish that the differential intrinsic properties of the PV and SST neurons, specifically the higher threshold and gain of the PV neurons, are in and of themselves sufficient to drive a distinct role of these inhibitory neurons classes in Up-state dynamics. While previous work has focused on the differential synaptic and connectivity patterns of PV and SST to cortical function, the current results emphasize that the differential threshold and gain of the F-I function of these inhibitory subclasses may underlie their distinct functional roles.

## Acknowledgments

We would like to thank Rodrigo Laje and Saray Soldado-Magraner for comments on earlier versions of this manuscript, and Sotiris Masmanidis, Carlos-Portera Cailliau, Peyman Golshani, Michael Seay for helpful scientific discussions and technical assistance and/or for their comments on earlier versions of this manuscript. This work was supported by the National Institute of Mental Health grant MH60163), and supplement MH060163-15S1.

## DATA and CODE AVAILABILITY

The simulation code and the analyses code and data used for the main results will be made available on https://github.com/BuonoLab/UpStates_2020

## AUTHOR CONTRIBUTIONS

J.L.R-S, H.M and D.V.B. conceived and designed the experiments; J.L.R-S performed most experiments with the help of H.M.; J.L.R-S analyzed the experimental data; D.V.B. performed the computational simulations and analyses; J.L.R-S prepared most figures; D.V.B. drafted manuscript and J.L.R-S and H.M revised the manuscript; J.L.R-S, H.M and D.V.B. approved the final version of the manuscript.

## COMPETING INTERESTS

The authors declare no competing interests.

---

**Extended Figure 7-1.**
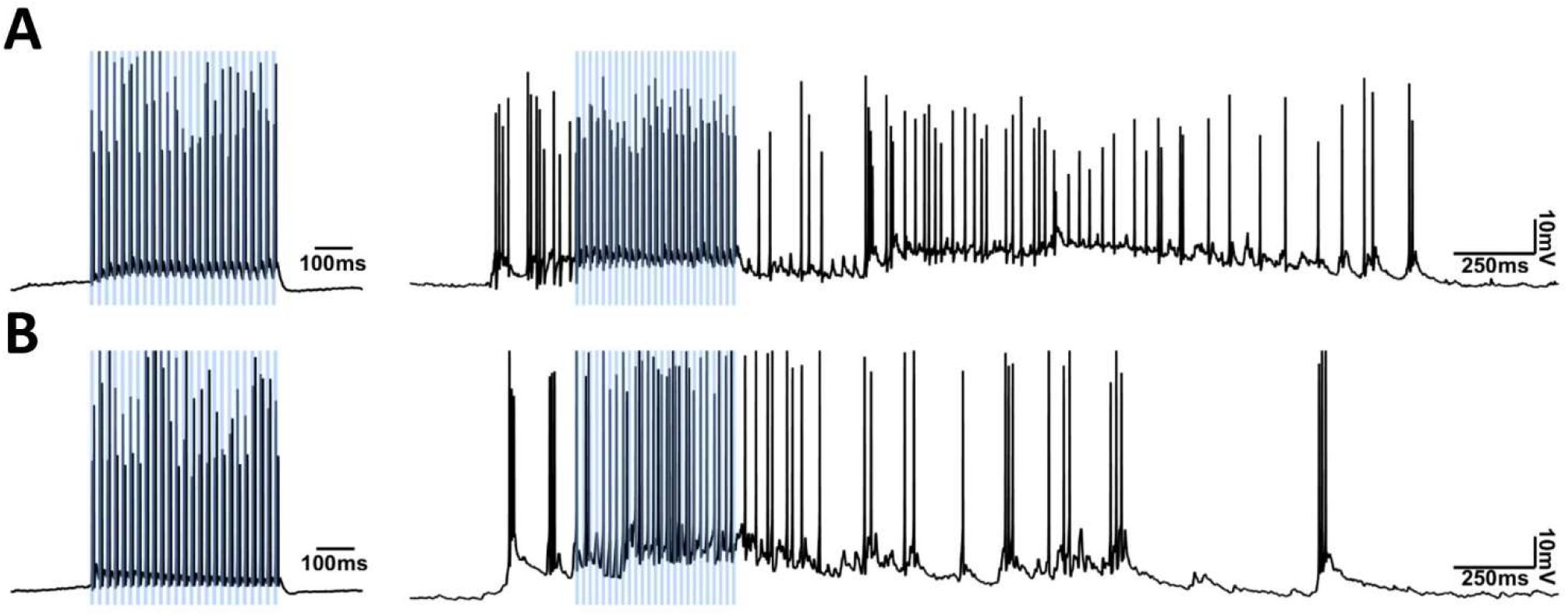
Light stimulation increases the firing rate of ChETA positive inhibitory neurons during Down- and Up-states. **(A)** Light stimulation depolarizes a PV ChETA-positive neuron during a Down-state (left) and an Up-state (right). **(B)** Same as A but with an SST neuron.

**Extended Figure 7-2.**
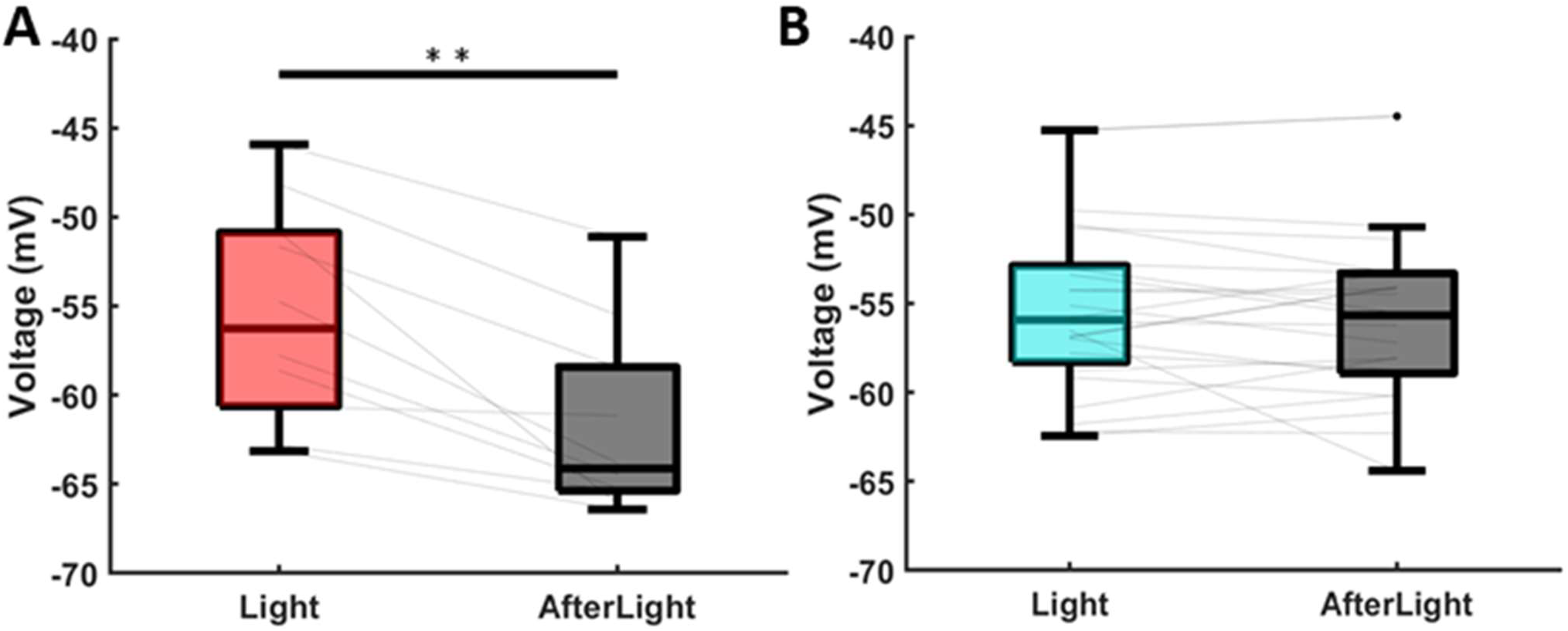
Activation of PV neurons reduces membrane potential during Up-States. **(A)** The median Pyr voltage was significantly reduced 100ms following light stimulation in PV neurons compared during the light stimulation. A much lower voltage signifies that a transition from an Up to Down state has happened. **(B)** Same as A but with an SST neuron light stimulation. There is no difference in the voltages.

## Notes

### Competing Interest Statement

The authors have declared no competing interest.

